# The memory of airway epithelium damage in smokers and COPD patients

**DOI:** 10.1101/2021.04.21.439437

**Authors:** François M. Carlier, Bruno Detry, Marylène Lecocq, Amandine M. Collin, Thomas Planté-Bordeneuve, Stijn E. Verleden, Monique Delos, Benoît Rondelet, Wim Janssens, Jérôme Ambroise, Bart M. Vanaudenaerde, Sophie Gohy, Charles Pilette

**Affiliations:** Pole of Pneumology, ENT, and Dermatology, Institute of Experimental and Clinical Research, Université Catholique de Louvain, Brussels, Belgium; Department of Pneumology, CHU Mont-Godinne UCL Namur, Yvoir, Belgium; Lung Transplant Centre, CHU Mont-Godinne UCL Namur, Yvoir, Belgium; Department of chronic diseases, metabolism and ageing, Katholieke Universiteit Leuven, Leuven, Belgium; Department of Pathology, CHU Mont-Godinne UCL Namur, Yvoir, Belgium; Deparment of Cardiovascular and Thoracic Surgery, CHU Mont-Godinne UCL Namur, Yvoir, Belgium; Centre de Technologies Moléculaires Appliquées, Institute of Experimental and Clinical Research, Université catholique de Louvain, Brussels, Belgium; Department of Pneumology, Cliniques Universitaires St-Luc, Brussels, Belgium; Cystic Fibrosis Reference center, Cliniques Universitaires St-Luc, Brussels, Belgium

**Keywords:** COPD, epithelial biology, barrier dysfunction, EMT, pIgR

## Abstract

**Rationale:** Chronic obstructive pulmonary disease (COPD) is a devastating and mostly irreversible lung disease. In COPD, the bronchial epithelium displays several structural and functional abnormalities affecting barrier integrity, cell polarity, and differentiation, as well as epithelial-to-mesenchymal transition and inflammation. Although COPD displays mostly irreversible changes, the (ir)reversible nature of epithelium pathology *ex vivo* remains poorly known and was the aim of this study.

**Methods:** The persistence of COPD epithelium abnormalities was addressed in long-term (10 weeks) primary cultures of air/liquid interface-reconstituted airway epithelium from non-smoker controls, smoker controls, and COPD patients. Barrier function, epithelial polarity, cell commitment, epithelial-to-mesenchymal transition and inflammation were assessed *in vitro*, and certain features were compared *in situ* to the native epithelium. The role of inflammation was explored by stimulating cultures with a cytokine mix consisting of TNF-α, IL-6 and IL-1β.

**Measurements and main results:** Almost all epithelial defects (barrier dysfunction, impaired polarity, lineage abnormalities) observed in cells from smokers and COPD patients persisted *in vitro* up to week 10, except IL-8/CXCL-8 release and epithelial-to-mesenchymal transition which declined over time. Cell lineage and polarity impairments matched abnormalities observed *in situ* in the surgical samples from which the *in vitro* epithelium was derived. Cytokine treatment induced COPD-like changes and, in COPD cells, reactivated epithelial-to-mesenchymal transition.

**Conclusions:** The airway epithelium from smokers and COPD patients displays a memory of its native state and previous injuries by cigarette smoking, which is multidimensional and sustained for extended periods of time.

## Introduction

Chronic obstructive pulmonary disease (COPD) constitutes the third cause of death worldwide (1), and is mainly due to noxious airborne stimuli, most importantly cigarette smoke (2). COPD is characterized by the irreversible narrowing and disappearance of small conducting airways (3) and destruction of alveolar walls (4). Current therapies provide limited clinical and functional benefits and fail at targeting the underlying pathways leading to structural remodeling of the lungs (5).

The airway epithelium (AE), as first line barrier against inhaled particles, is constantly exposed to airborne pollutants, and fulfils multiple functions to maintain pulmonary homeostasis. It ensures adequate barrier function, cell differentiation and polarization, while maintaining a tight control on inflammatory mechanisms. In COPD, the AE fails at these duties, exhibiting altered physical barrier function (6, 7) underpinned by decreased expression of apical junction complexes (AJCs) proteins (8), abnormal differentiation and function of ciliated and club cells (9–12) as well as hyperplastic goblet cells (13), and aberrant epithelial-to-mesenchymal transition (EMT) that is triggered by cigarette smoke (14) and further enhanced in COPD (15). In addition, the polymeric immunoglobulin receptor (pIgR)/secretory component (SC) system, which ensures the transcytosis and release of polymeric immunoglobulins into mucosal secretions, witnessing epithelial polarization (16, 17) is also defective in COPD, with decreased epithelial pIgR expression (16) and local S-IgA deficiency in small airways (18, 19). Finally, epithelial inflammation control is dysregulated in COPD (20), with increased intraepithelial neutrophils (21) and sputum tumor necrosis factor (TNF)-α, interleukin-8 (IL-8/CXCL-8) and neutrophils (22, 23).

The air/liquid interface (ALI) culture model of primary human bronchial epithelial cells (HBEC) allows the reconstitution of the AE *in vitro* and recapitulates several of the alterations observed *in situ* in COPD. Cigarette smoke-exposed COPD HBEC show decreased E-cadherin and ZO-1 expression as well as reduced transepithelial electric resistance (TEER) when compared with smokers without COPD (14, 24) and after 4 weeks culture, the COPD ALI-AE displays altered lineage differentiation, decreased pIgR expression, and EMT (8, 10, 16), along with increased cytokine release (25). Although these data suggest the persistence of AE abnormalities, it remains unclear whether (and to what extent) these structural changes persist on the long-term, matching the irreversible nature of COPD. COPD patients who quit smoking for more than 3.5 years display less goblet cell hyperplasia (26) and smoking cessation may improve lung function and survival (27, 28). Conversely, smoking cessation does not influence COPD-related epidermal growth factor receptor activation (26) or protease activity (29). In COPD, the inflammatory pattern shared with ‘healthy’ smokers is amplified and persists even after smoking cessation, although with conflicting data (27, 30, 31).

Whereas small airway changes are importantly involved in COPD pathophysiology, the recent *Lancet Commission* advocates for a broader definition including symptoms related to large airways (e.g., bronchitis) and for active research in the field (32). The present study aims to elucidate whether epithelial changes in large airways are persistently imprinted, questioning the disease memory (31) retained in the COPD epithelium. To address this question, we reconstituted HBEC-derived ALI-AE from non-smokers, non-COPD smokers and COPD patients, and cultured it *in vitro* for up to 10 weeks. The spontaneous evolution of abnormalities was assayed for epithelial readouts including barrier function, cell differentiation, EMT, pIgR/SC-related polarity and production of inflammatory cytokines. In addition, we assessed whether exogenous inflammation could trigger COPD-related changes.

## Materials and methods

### Study population and lung tissue samples

Lung surgical specimens were obtained from patients with or without COPD. Subjects were sorted on basis of clinical diagnosis and pulmonary function tests, according to the GOLD 2001 classification (33). A primary proximal AE was reconstituted *in vitro* from all subjects and subjected to long-term culture (10 weeks, n=26) or mid-term culture with cytokine stimulation (5 weeks, n=25). Tables 1 and S1 recapitulate patients’ characteristics for each population. Detailed information of methods is available in the online data supplement.

### *In vitro* reconstitution of primary human AE on ALI culture

ALI-cultures were conducted as previously published (17). TEER was assessed every week for each sample, using the EMD Millipore™ Millicell-ERS Volt-Ohmmeter (Fisher Scientific, Hampton, NH, USA) after transiently filling the apical pole with sterile PBS and correcting for the resistance of the transwell membrane.

### Reverse transcriptase quantitative polymerase chain reaction (RT-qPCR)

RNA extraction, reverse transcription, and RT-qPCR were performed as previously described (34).

### Western blot assays

HBEC lysates were analyzed by Western blot as previously described (35) except for revelation, performed by chemiluminescence (GE Healthcare, Pittsburgh, PA) before detection by Chemidoc XRS apparatus (Bio-Rad) and quantification by Quantity One software (Bio-Rad).

### Enzyme-linked immunosorbent assay (ELISA) for SC, IL-8/CXCL-8, IL-6, and fibronectin

Levels of SC (in apical washes) and cytokines (in basolateral supernatants) were measured by sandwich ELISA, as previously described (11, 35). Fibronectin release in the basal medium was assessed by direct ELISA as previously reported (8, 36).

### Immunofluorescence staining using tyramide signal amplification

Five micron-sections of surgical lung tissue or reconstituted ALI epithelium were prepared and stained as previously described (17). Detailed information of methods and quantification is provided in the online data supplement.

### Statistical analysis

Data were expressed as means and standard deviation when passing normality tests, otherwise as medians and interquartile ranges, and were analyzed with JMP® Pro, Version 14 (SAS Institute Inc., Cary, NC, USA) and GraphPad Prism version 8.0.2 for Windows (GraphPad Software, La Jolla, CA, USA). p-values < 0.05 were considered statistically significant. Detailed information of statistical analysis is provided in the online data supplement.

### Results

Readouts were assessed (for each sample, according to availability) at different timepoints of redifferentiation, referred to as early (1 week ALI), short-term (2-3 weeks), mid-term (4-7 weeks) and long-term (8-10 weeks) cultures.

### Barrier and junctional properties in the COPD AE

The AE physical barrier was assessed by measuring TEER of ALI-AE up to 10 weeks. According to smoking status and lung function, the study population was divided into 4 groups: non-smoker controls (NS; n=5), smoker controls (Smo, n=7), mild/moderate COPD (COPD1-2; n=7) and severe/very severe COPD (COPD3-4; n=7). The characteristics of the study population are summarized in Table 1.

**Table 1.**
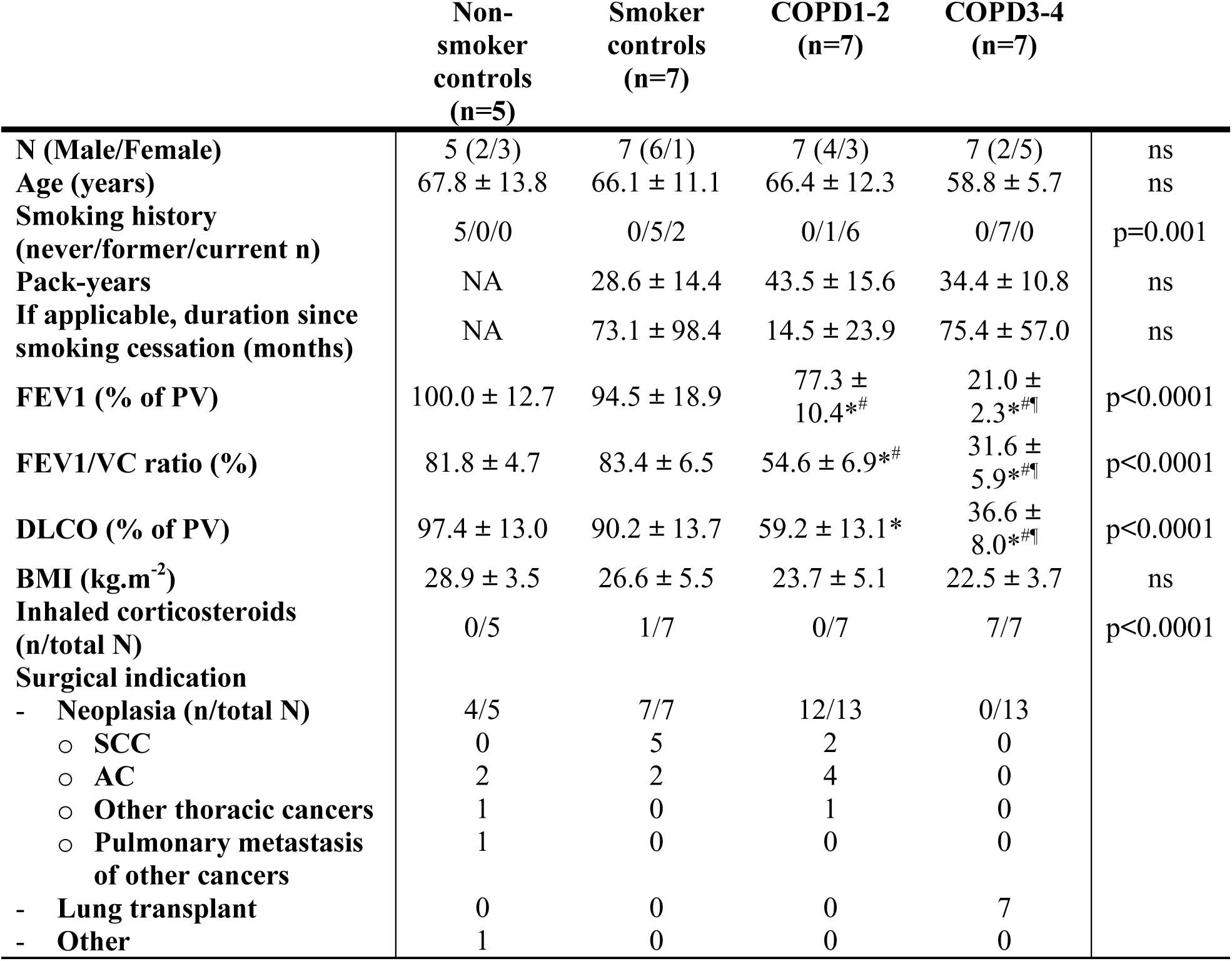
Patient cohort for ALI-cultures. Data are presented as mean ± SD, unless otherwise stated. Demographic data, lung function tests, smoking history and inhaled corticotherapy are stated for the patient groups, classified according to smoking history and the presence and severity of airflow limitation. AC, adenocarcinoma; ALI, air/liquid interface; BMI, body mass index; COPD, chronic obstructive pulmonary disease; DLCO, diffusing capacity of the lung for CO; FEV1, forced expiratory volume in 1 s; PV, predicted values; SCC, squamous cell cancer; SD, standard deviation; VC, vital capacity. * = p<0.05 compared to non-smoker controls ^#^ = p<0.05 compared to smoker controls ^¶^ = p<0.05 compared to COPD stage 1-2 patients ns, not significant

The COPD AE displayed decreased TEER as compared with NS, and to a lesser extent with Smo, which also showed decreased TEER compared with NS (Figure 1a). This defect appeared in early cultures, persisting in long-term cultures (Figure 1b). In addition, TEER inversely correlated with the disease severity witnessed by the forced expiratory volume in one second (FEV1). This correlation was significant from early up to long-term cultures (Figure 1c).

**Figure 1.**
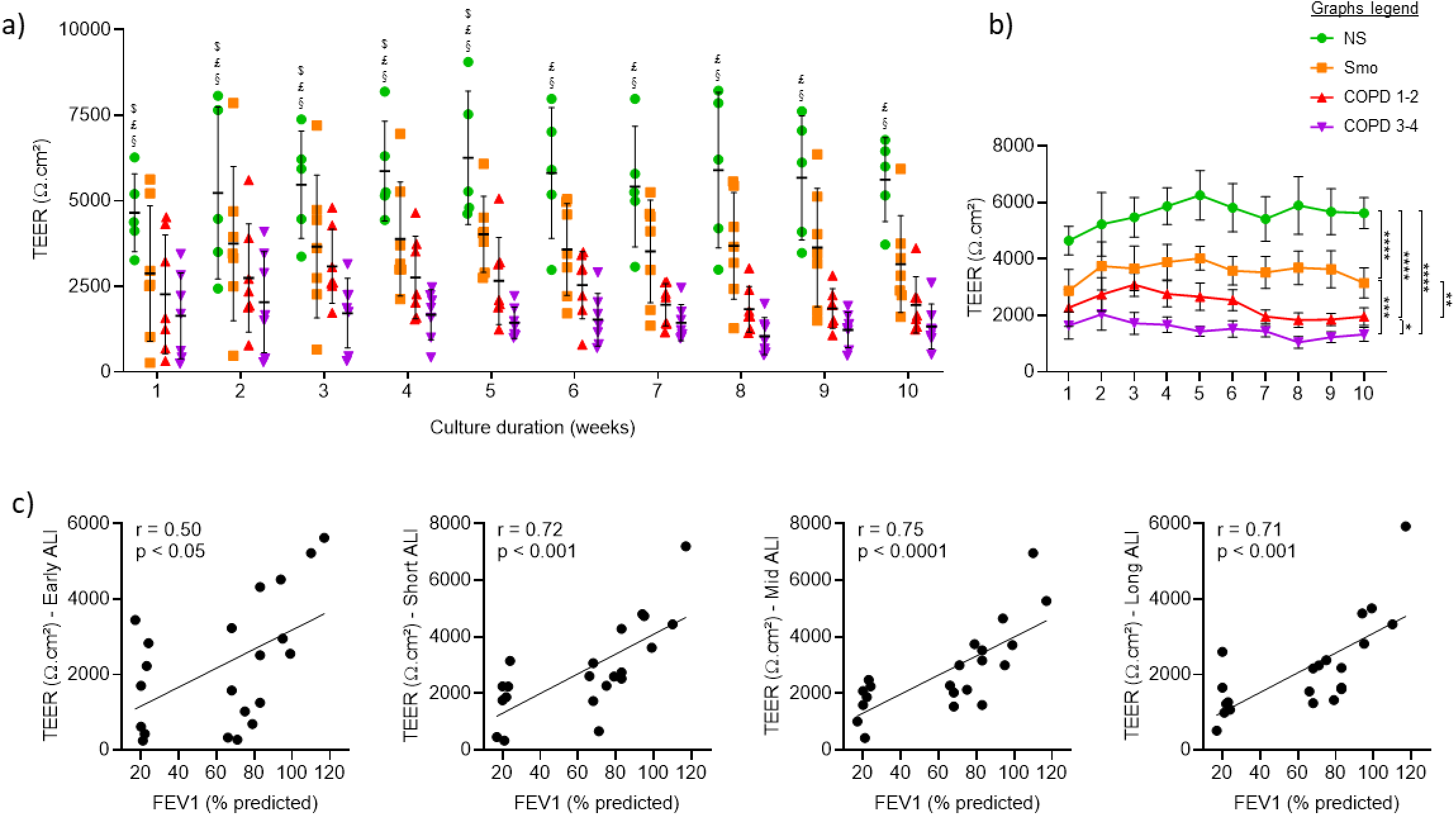
COPD and Smo AE displays persistently decreased TEER compared with NS AE. a) TEER in the ALI-AE from NS, Smo and COPD patients (GOLD classification from 1 to 4 according to the spirometric severity of the disease). ALI-AE derived from COPD patients display a sharp decrease in TEER as compared with (non)-smokers at all time periods, that is also observed to a lesser extent in mild and moderate COPD. $, £ and § highlight significant decreases in Smo, COPD1-2 and COPD3-4 patients respectively, as compared to NS. b) Longitudinal analysis of the evolution of the TEER in the ALI-AE from NS, Smo, mild-to-moderate COPD and (very) severe COPD patients, showing a smoking-related persistent barrier dysfunction that is further enhanced in COPD. c) The barrier dysfunction observed in COPD, witnessed by the TEER decrease, significantly correlates with the disease severity assessed by the FEV1, from early cultures up to long-term cultures. Graphs include only Smo and COPD samples to specifically assess the correlation with the disease severity. ***, **** indicate p-values of less than 0.001, and 0.0001, respectively. Bars indicate mean ± standard deviation. AE, airway epithelium; ALI, air-liquid interface; COPD, chronic obstructive pulmonary disease; FEV1, forced expired volume in 1 second; NS, non-smokers; SEM, standard error of the mean; Smo, smokers; TEER, transepithelial electric resistance.

The molecular substratum of this long-lasting barrier disruption was questioned by assessing mRNA abundance and protein expression of major components of AJCs, namely claudin-1 (*CLDN1*), E-cadherin (*CDH1*), occludin (*OCLN*) and ZO-1/tight junction protein 1 (*TJP1*). Although no difference was observed regarding mRNA (Figure S1), protein expression of E-cadherin was reduced in COPD versus NS in early and short-term ALI (Figure 2a), while that of occludin was decreased from early up to long-term cultures in Smo and COPD (Figure 2b), and inversely correlated with FEV1 up to mid-term (Figure 2c). Figure 2d shows representative blots for E-cadherin and occludin in NS and COPD3-4 AE.

**Figure 2.**
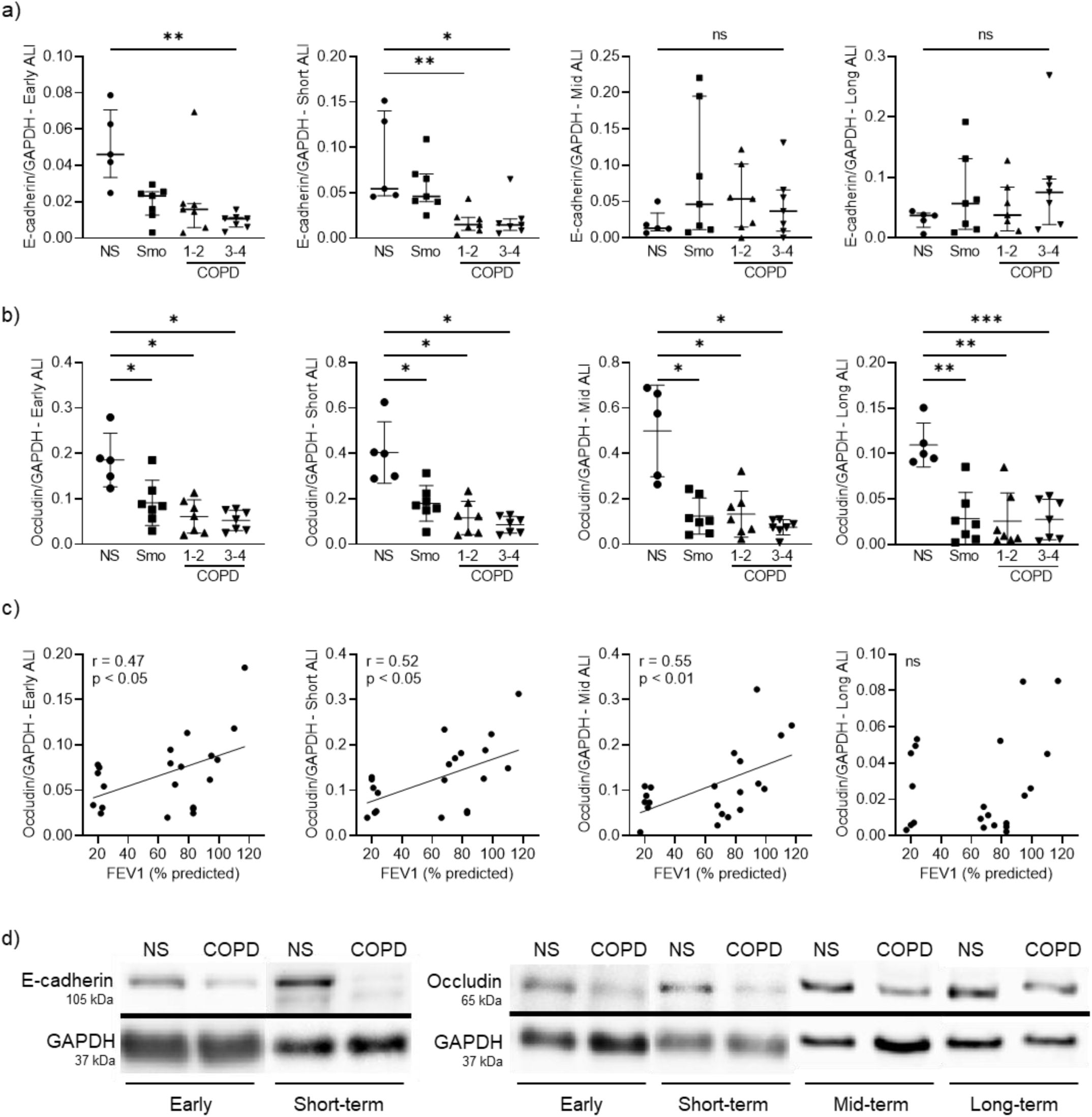
COPD and Smo AE displays decreased E-cadherin and occludin protein expression compared with NS. a) Decreased E-cadherin protein levels in early and short-term ALI-AE, but no more at mid-term and long-term, in Smo and COPD ALI-AE as compared with that of NS. b) Decreased occludin protein levels, observable from early up to long-term cultures, in Smo and COPD ALI-AE as compared with that of NS. c) Occludin decrease observed in COPD, significantly correlates with the disease severity assessed by the FEV1, from early cultures up to mid-term (but not long-term) cultures. Graphs include only Smo and COPD samples to specifically assess the correlation with the disease severity. d) Representative blots for E-cadherin and occludin in NS and very severe COPD AE, showing decrease in E-cadherin expression in early and short-term ALI, and decrease in occludin expression in COPD ALI-AE from early up to long-term cultures. *, **, *** indicate p-values of less than 0.05, 0.01, and 0.001, respectively (analyzed using the Kruskal-Wallis test followed by Dunn’s post-hoc test). Bars indicate median ± interquartile range. AE, airway epithelium; ALI, air-liquid interface; COPD, chronic obstructive pulmonary disease; FEV1, forced expired volume in 1 second; NS, non-smokers; ns, not significant; SEM, standard error of the mean; Smo, smokers.

These data globally depict epithelial barrier dysfunction that is engaged in Smo, further worsens in COPD, and persists upon prolonged culture.

### Lineage differentiation of the COPD AE

The persistence of differentiation alterations in COPD was questioned by assessing specific markers and transcription factors of early differentiation towards intermediate cells, as well as of goblet and ciliated cells.

#### Epithelial pre-differentiation

mRNA abundance of MYB Proto-Oncogene (*MYB*), a marker of early differentiation of basal cells (37), was decreased up to long-term in Smo and COPD-derived AE (Figures S2a), correlating with FEV1 at some time-periods (Figure S2b).

#### Differentiation towards ciliated cells

The differentiation towards ciliated cells was assessed by measuring the mRNA abundance of Forkhead Box J1 (*FOXJ1*), a transcription factor involved in ciliated cells’ commitment, and of the dynein axonemal intermediate chain-1 (*DNAI1*), a marker of ciliated cells’ terminal differentiation. We also counted β-tubulin IV^+^ cells in ALI-AE and in native tissues. Smo-and COPD-derived cultures displayed persisting decreases of both gene transcripts, which correlated with FEV1 at some time-periods (Figures 3a-b & S3a-b). β-tubulin IV^+^ (ciliated) cells numbers were decreased up to long-term ALI-cultures from Smo and COPD (Figures 3c-d), in line with reduced β-tubulin IV^+^ surface in the native COPD AE (Figure 3e).

**Figure 3.**
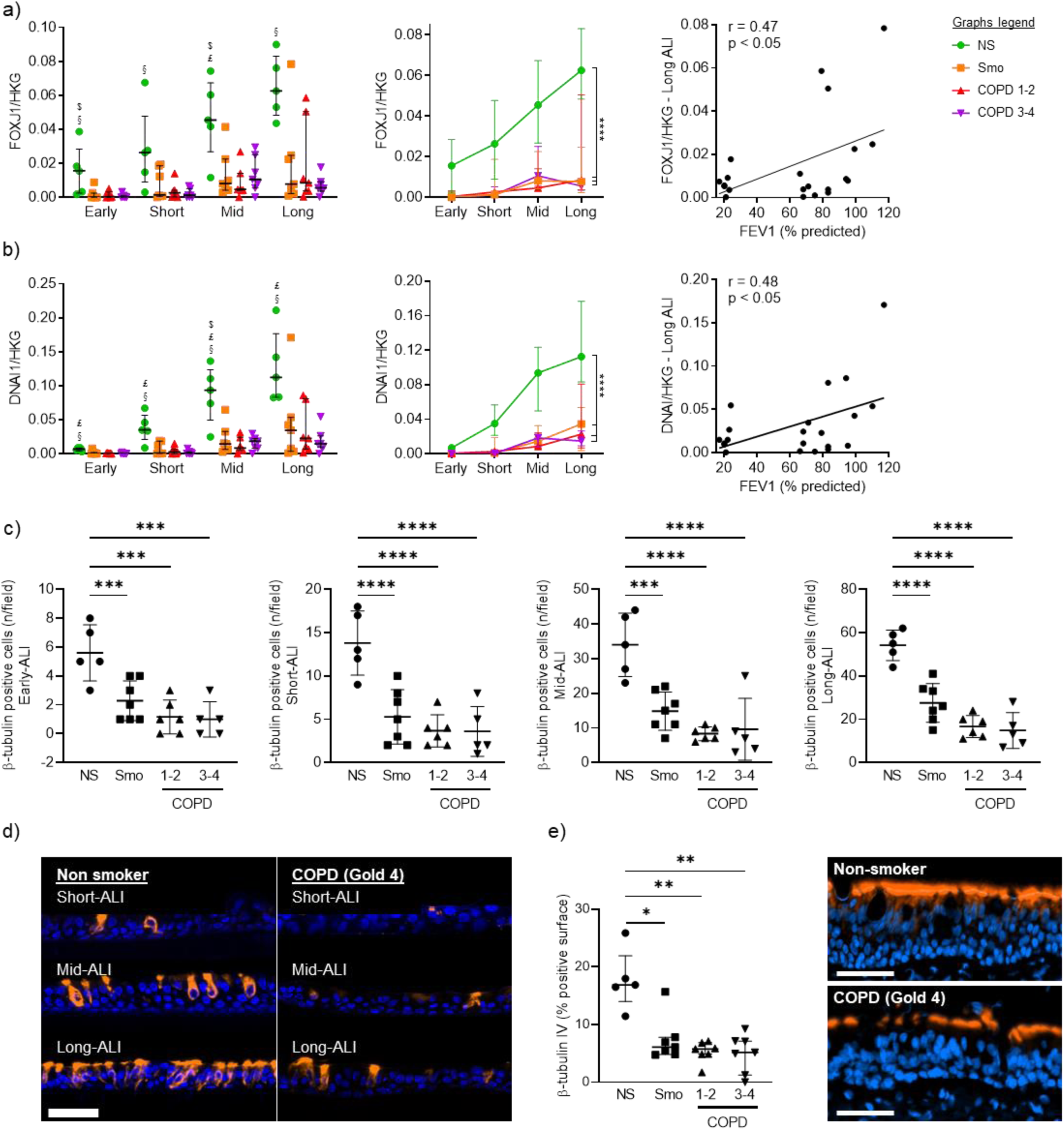
Altered differentiation programming towards ciliated cells in Smo and COPD patients. a) Decreased *FOXJ1* expression in Smo and COPD AE, from early up to long-term ALI culture. Longitudinal analysis (middle graph) shows a strong, persistent downregulation in Smo and COPD, as compared with NS. At some time periods (as represented here in long-term ALI-AE), *FOXJ1* expression correlated with the severity of the disease witnessed by the FEV1 (left graph). Correlation graph includes only Smo and COPD samples to specifically assess the correlation with the disease severity. $, £ and § highlight significant decreases in Smo, COPD1-2 and COPD3-4 patients respectively, as compared to NS. b) Decreased *DNAI1* expression in Smo and COPD AE, from early up to long-term ALI culture. Longitudinal analysis (middle graph) shows a strong, persistent downregulation in Smo and COPD, as compared with NS. At some time periods (as represented here in long-term ALI-AE), *DNAI1* expression correlated with the severity of the disease witnessed by the FEV1 (left graph). Correlation graph includes only Smo and COPD samples to specifically assess the correlation with the disease severity. c) Decreased numbers of bêta-tubulin IV positive ciliated cells in Smo and COPD-derived ALI at every time-period. d) Illustrative TSA-enhanced immunofluorescence staining for bêta-tubulin IV, demonstrating the differentiated emergence of ciliated cells in NS and COPD-derived ALI cultures. Scale bar, 50µm. e) Quantification of bêta-tubulin IV^+^ surface in bronchial sections demonstrating reduced *in situ* expression of bêta-tubulin IV in native Smo and COPD AE, as illustrated by TSA-enhanced immunofluorescence staining for bêta-tubulin IV (right picture; scale bar, 50µm.). *, **, ***, **** indicate p-values of less than 0.05, 0.01, 0.001, and 0.0001, respectively. Bars indicate median ± interquartile range (panels a, b, and e) or mean ± standard deviation (panel c). AE, airway epithelium; ALI, air-liquid interface; COPD, chronic obstructive pulmonary disease; CT, control; FEV1, forced expired volume in 1 second; HKG, housekeeping genes; NS, non-smokers; SEM, standard error of the mean; Smo, smokers; y, years.

#### Differentiation towards goblet cells

The mRNA abundance of SAM Pointed Domain Containing ETS Transcription Factor (*SPDEF*) and Forkhead Box A3 (*FOXA3*) were assessed as transcription factors inducing goblet cell differentiation (37–39), along with the protein expression of MUC5AC, a terminal product of goblet cell differentiation. Compared to NS, *SPDEF* expression was persistently increased in Smo as well as in COPD, although to a lesser extent (Figure 4a). Similarly, MUC5AC^+^ goblet cells numbers were increased in Smo ALI-cultures versus NS (Figure 4b), matching *in situ* findings, where MUC5AC^+^ surface in Smo-AE was considerably larger than in NS (Figure 4c). As for *SPDEF*, *FOXA3* was upregulated in Smo compared with NS (Figure S3c).

**Figure 4.**
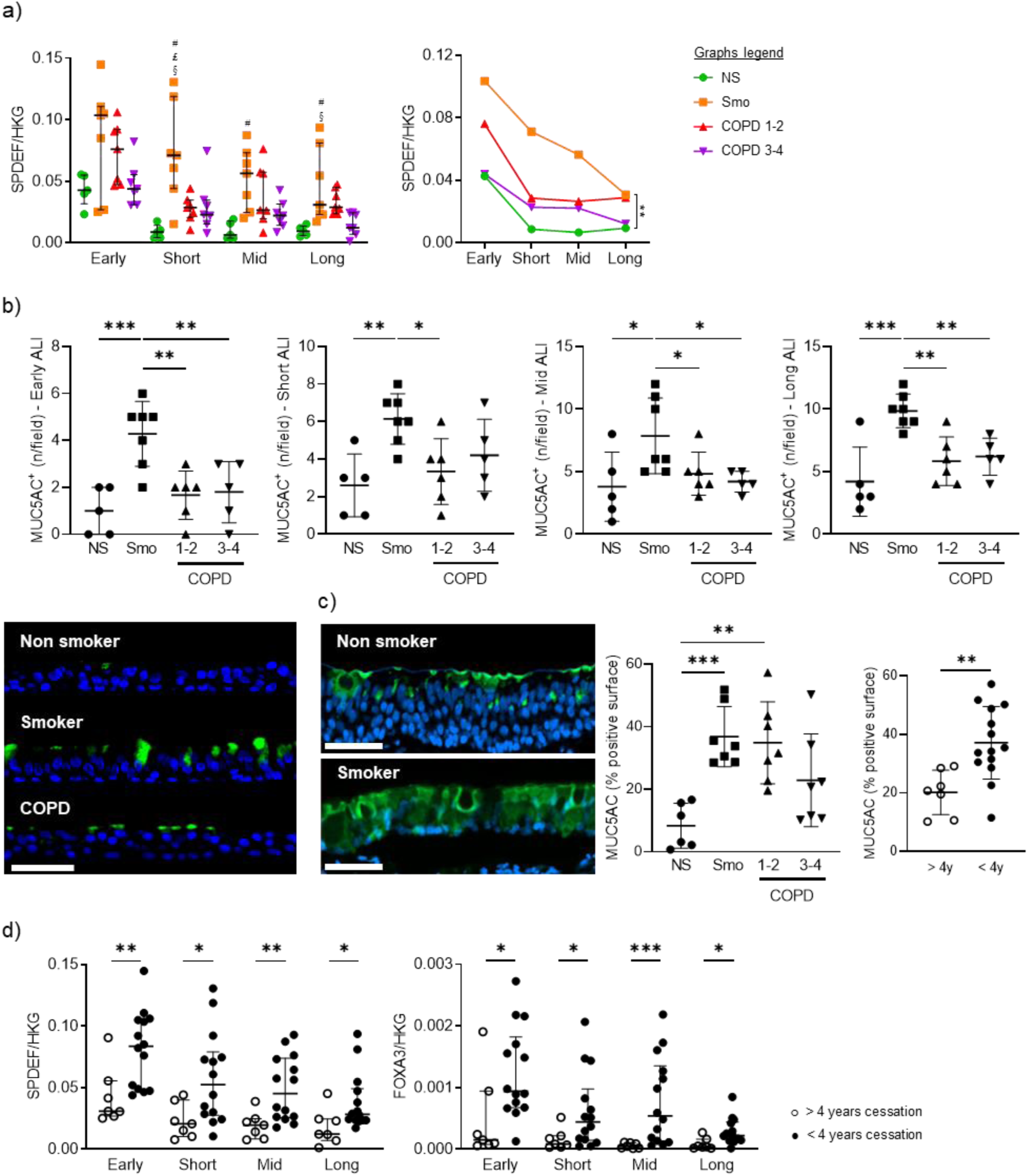
Altered differentiation programming towards goblet cells in Smo and COPD patients. a) Increased *SPDEF* expression in Smo AE as compared with NS in short-term, mid-term and long-term ALI culture, as depicted for each timepoint (left graph) and in a longitudinal analysis (right graph, significance bar: Smo *versus* NS). Of note, error bars were not added on the longitudinal data for reasons of readability. #, £ and § highlight significant decreases in NS, COPD1-2, and COPD3-4 patients respectively, as compared to NS. b) Increased MUC5AC positive goblet cells in Smo AE as compared with NS and COPD patients in early, short-term, mid-term and long-term ALI culture. Bottom left picture: illustrative TSA immunofluorescence staining for MUC5AC, showing increased positive goblet cells in ALI cultures (here, in mid-term ALI-AE) from Smo as compared with NS and COPD. c) *In situ*, in bronchial surgical sections, MUC5AC positive surface is increased in Smo AE as compared with NS and COPD. In addition, MUC5A positive surface is significantly higher in patients who quit smoking for less than 4 years (plain dots) as compared with long-quitters (hollow dots). d) At every time period, *SPDEF* (left graph) and FOXA3 (right graph) mRNA levels are decreased in ALI-AE from Smo who quit smoking for more than 4 years (plain dots) as compared with long quitters (hollow dots). *, **, *** indicate p-values of less than 0.05, 0.01, 0.001, and 0.0001. Bars indicate means ± standard deviation (panels a, b, and c) or medians ± interquartile ranges (panel d). AE, airway epithelium; ALI, air-liquid interface; COPD, chronic obstructive pulmonary disease; CT, control; FEV1, forced expired volume in 1 second; HKG, housekeeping genes; NS, non-smokers; SEM, standard error of the mean; Smo, smokers; y, years. Scale bar, 50µm.

Both *SPDEF* and *FOXA3* expression were decreased at all timepoints in smokers who quit smoking for more than 4 years, versus active smokers and ex-smokers with a smoking cessation of less than 4 years (Figure 4d). These results corroborated *in situ* findings, with reduced MUC5AC^+^ surface in long-quitters’ AE (Figure 4c, right graph).

These data show that goblet cell hyperplasia relates to active/recent smoking and demonstrate its persistence over time *in vitro*.

### EMT of the COPD AE

The expression of EMT protein markers (vimentin, fibronectin) was assessed in long-term ALI cultures. Increased vimentin contents were observed in early, short-term and mid-term, but not long-term COPD3-4 versus NS cultures, as well as in early Smo- and COPD1-2 cultures (Figure 5a-b). Accordingly, COPD and Smo-AE displayed increased fibronectin release up to mid-term cultures versus NS (Figure 5c). No difference remained in long-term cultures, while no difference was observed regarding vimentin mRNA abundance (*VIM*, Figure S4). These data show that EMT features progressively vanish in COPD.

**Figure 5.**
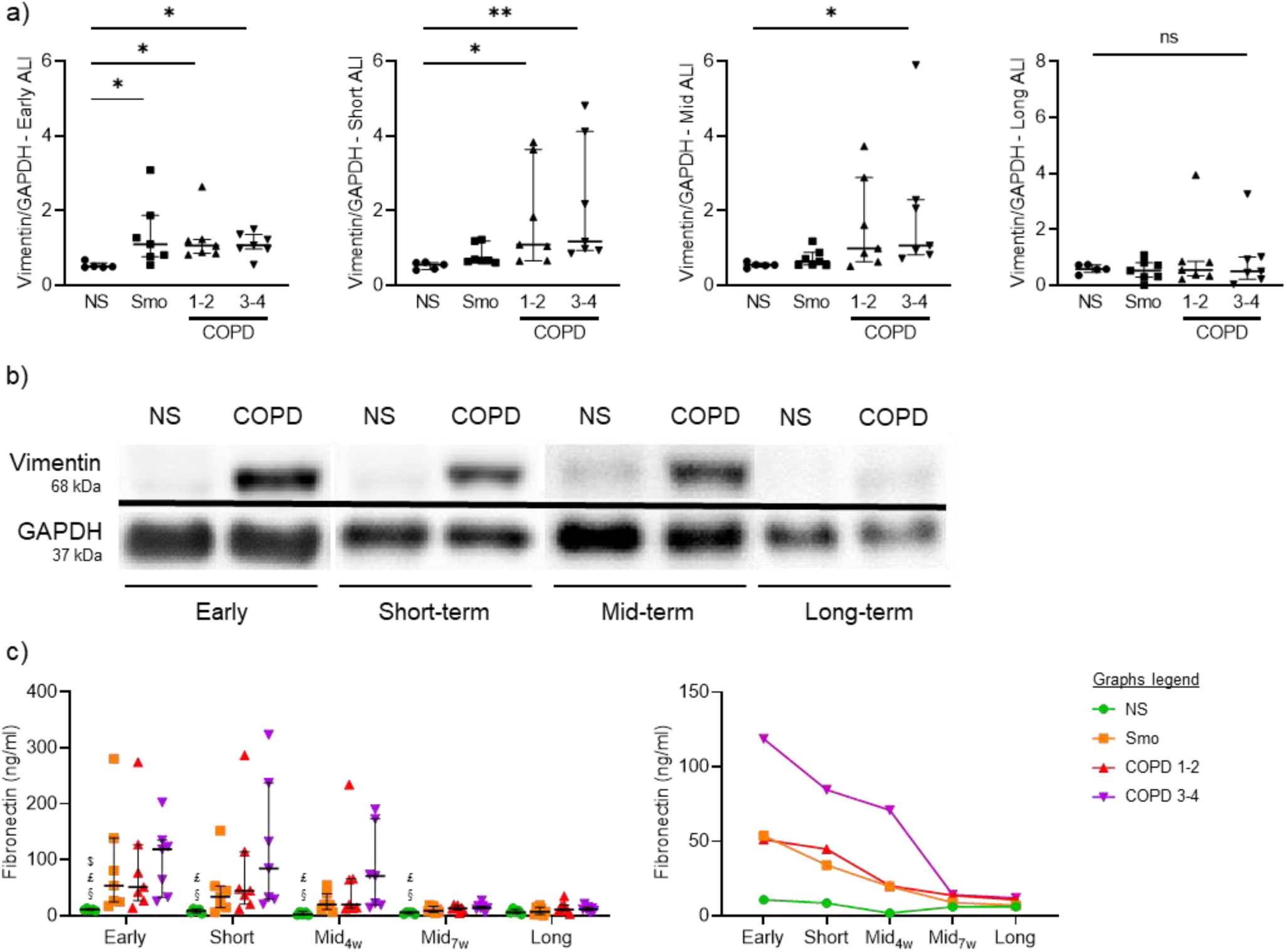
COPD-related EMT features vanish during long-term ALI cultures. a) Vimentin contents are increased in early Smo cultures, and up to mid-term in COPD ALI-AE, with this increase vanishing in long-term cultures. b) Illustrative gels from NS and very severe COPD-derived AE cultured at the ALI from early up to long-term, illustrating the vimentin increased contents up to mid-term in COPD. c) Increased fibronectin release disappears in mid-term cultures from Smo and mild-to-moderate COPD patients, while it persists up to later (7 weeks) in (very) severe COPD AE. *, ** indicate p-values of less than 0.05 and 0.01, respectively. Bars indicate median ± interquartile range. $, £, and § highlight significant decreases in Smo, COPD1-2 and COPD3-4 patients respectively, as compared to NS. AE, airway epithelium; ALI, air-liquid interface; COPD, chronic obstructive pulmonary disease; GAPDH, glyceraldehyde-3-phosphate dehydrogenase; NS, non-smokers; ns, not significant; Smo, smokers; w, weeks.

### Polarity-related pIgR/SC expression

Smo- and COPD-derived AE released less apical SC than NS, from short-up to long-term ALI cultures (Figure 6a), that was more prominent in COPD3-4 patients and correlated with disease severity (Figure 6b). Concordantly, COPD3-4 AE displayed decreased *PIGR* mRNA abundance throughout the long-term cultures. Although this was not significant on isolated time points (Figure 6c), longitudinal data demonstrated decreased *PIGR* mRNA abundance in COPD3-4 AE versus other groups (Figure 6c, right panel). In addition, considering that epithelial differentiation in ALI is completed at 5 weeks, the acquisition of maximal pIgR protein levels was delayed in COPD AE (Figure 6d). Concordantly, decreased pIgR levels were also observed *in situ*, in bronchial sections from Smo and COPD AE (Figure 6e).

**Figure 6.**
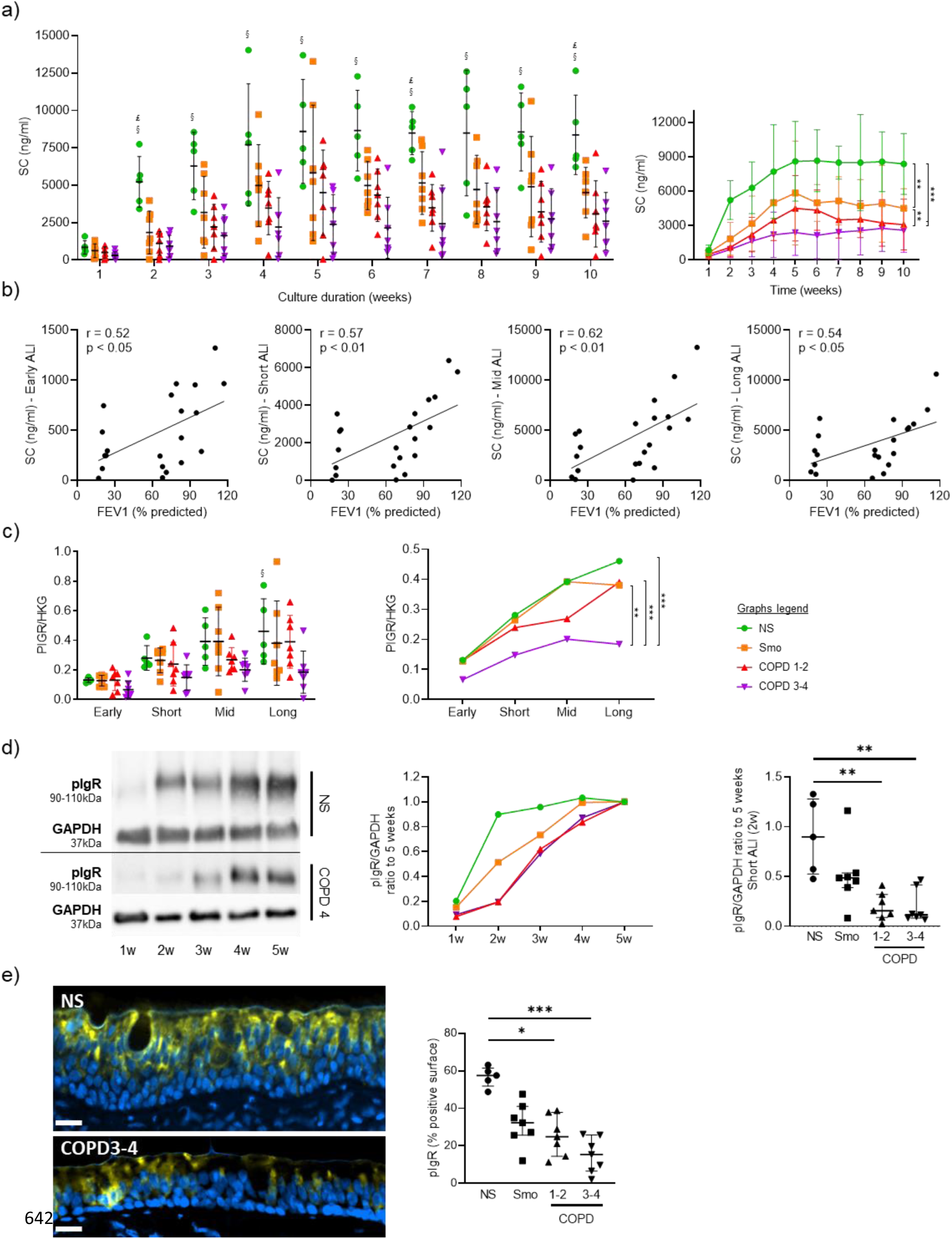
Impaired polarity, witnessed by a disruption and a delayed acquisition of PIGR/SC expression, is imprinted in the COPD AE. a) SC apical release in ALI-AE from Smo and COPD patients is decreased as compared with NS, from short-term ALI-AE up to long-term ALI-AE. $, £ and § highlight significant decreases in Smo, COPD1-2 and COPD3-4 patients respectively, as compared to NS. b) SC decreased release in the COPD AE moderately correlates with the disease severity, witnessed by the FEV1. Correlation graphs include only Smo and COPD samples to specifically assess the correlation with the disease severity. c) *PIGR* mRNA abundance is not significantly decreased in (very) severe COPD ALI-AE at separate time-points (left panel), but longitudinal analysis shows a significant decrease in *PIGR* expression in (very) severe COPD AE as compared with NS, Smo, and mild-to-moderate COPD (right panel). d) pIgR acquisition is delayed during the differentiation of the AE in COPD as compared with NS, as shown in the representative blots from NS and very severe COPD ALI-AE and summarized in the middle panel. The gap was maximal at 2 weeks ALI culture (right panel) and catches it up only after 4 weeks. e) TSA immunofluorescence staining for pIgR, showing reduced positive surface *in situ* (i.e., in bronchial sections) in the COPD AE as compared with that of NS. *, **, *** indicate p-values of less than 0.05, 0.01, and 0.001, respectively (analyzed using the Kruskal-Wallis test followed by Dunn’s post-hoc test, except for longitudinal analysis in B and C, mixed model). Bars indicate mean ± SD (a, c [left], d [right], e), dots indicate mean (c [right], d [left]). AE, airway epithelium; ALI, air-liquid interface; COPD, chronic obstructive pulmonary disease; FEV1, forced expired volume in 1 second; GAPDH, glyceraldehyde-3-phosphate dehydrogenase; HKG, housekeeping genes; NS, non-smokers; ns, not significant; pIgR, polymeric immunoglobulin receptor; PV, predicted values; SC, secretory component; SD, standard deviation; Smo, smokers; w, weeks.

These data show that both pIgR expression and functionality (SC release) are impaired in COPD. Along with impaired AJCs (see above), this indicates that the polarity of the AE in COPD is persistently altered.

### Inflammatory cytokine production by the COPD AE

IL-8/CXCL-8 and IL-6 are two main epithelial pro-inflammatory cytokines and were assayed in ALI-AE. A trend for increased *CXCL8* mRNA abundance was seen in Smo and COPD, in early to mid-term culture (Figure S5a). Accordingly, IL-8/CXCL-8 release was strongly increased up to mid-term cultures from Smo- and COPD-derived versus NS, whilst this difference disappeared afterwards (Figure 7a). Although no difference was observed regarding *IL6* mRNA abundance (Figure S5b), IL-6 production was increased in Smo- and COPD -AE. In contrast to IL-8/CXCL-8, IL-6 upregulation persisted in Smo and COPD long-term cultures (Figure 7b).

**Figure 7.**
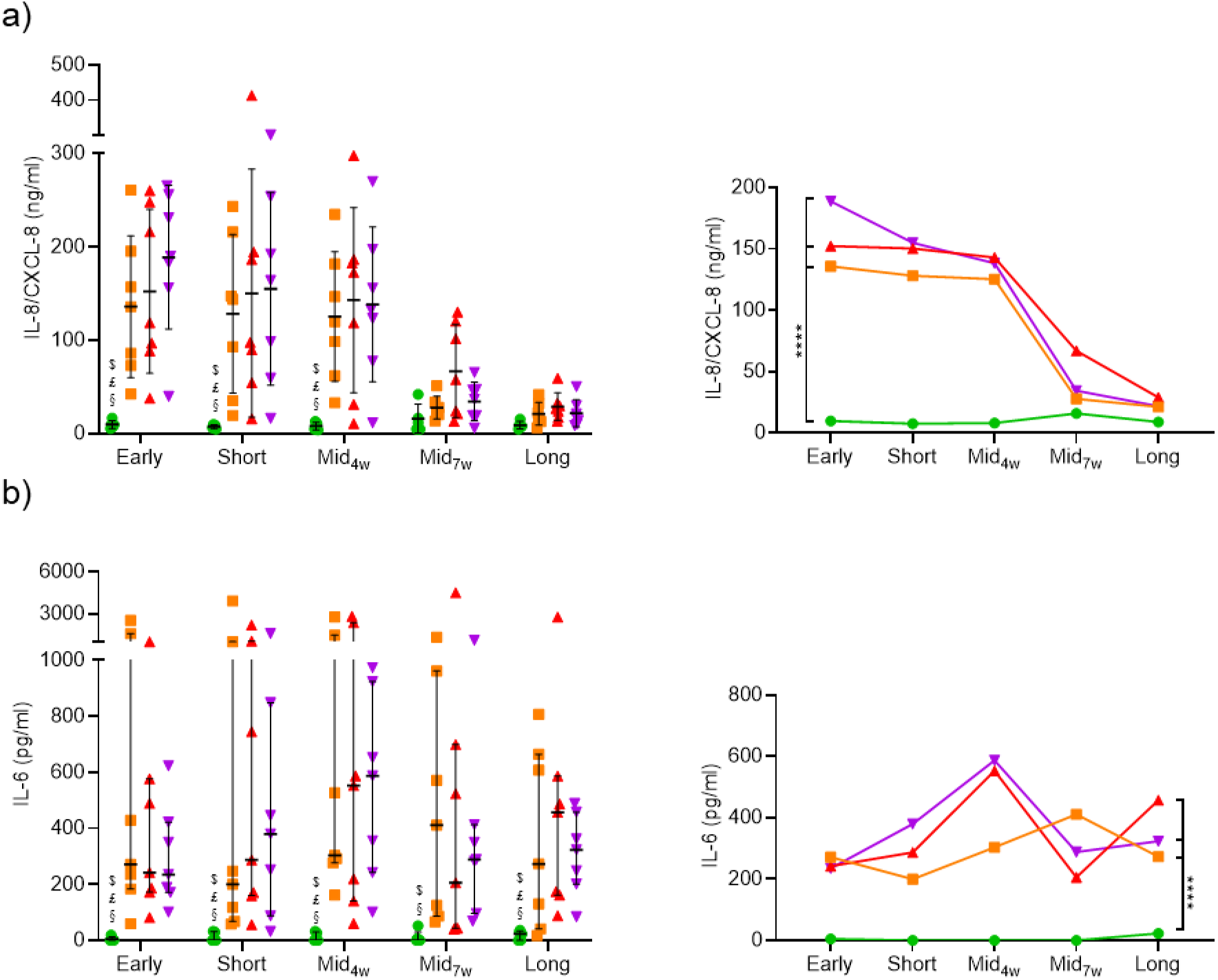
Epithelial release of inflammatory cytokines partly persists in COPD AE. a) IL-8/CXCL-8 production by the reconstituted ALI-AE is increased in early, short-term and mid-term ALI-AE from and COPD patients, but disappears afterwards, although a non-significant upward trend persists. b) IL-6 production by the reconstituted ALI-AE is increased in Smo and COPD and persists in long-term cultures. **** indicates p-values of less 0.0001. Bars indicate mean ± SD (panel a) and median ± interquartile range (panel b). AE, airway epithelium; ALI, air-liquid interface; COPD, chronic obstructive pulmonary disease; IL, interleukin; NS, non-smokers; ns, not significant; SD, standard deviation; Smo, smokers; w, weeks.

### Exogenous inflammation induces COPD-like AE features

Whether exogenous inflammation could promote COPD-like changes in the AE was assessed by supplementing the culture medium with IL-6, TNF-α and IL-1β (all at 5ng/ml) for up to 5 weeks. First, TEER was decreased in inflammatory conditions in early cultures with this defect persisting in subsequent weeks (Figure 8a). Cytokine-exposed cultures displayed similar values irrespectively of the original phenotype, possibly indicating a maximal effect on barrier (dys)function at the used concentrations. Second, cultures exposed to inflammatory cytokines displayed EMT features, with increased fibronectin release and vimentin contents (Figure 8b-c). Interestingly, this induction that was not significant in NS, was strikingly increased in COPD compared to pooled controls. These results show that the COPD AE, whilst losing its intrinsic EMT features, remains prone to develop EMT upon inflammatory exposure. Third, cytokine-induced inflammation altered epithelial polarity, with SC apical release being decreased in all cultures except in COPD3-4, probably due to pre-existent pIgR/SC severe impairment in this group (Figure 8d).

**Figure 8.**
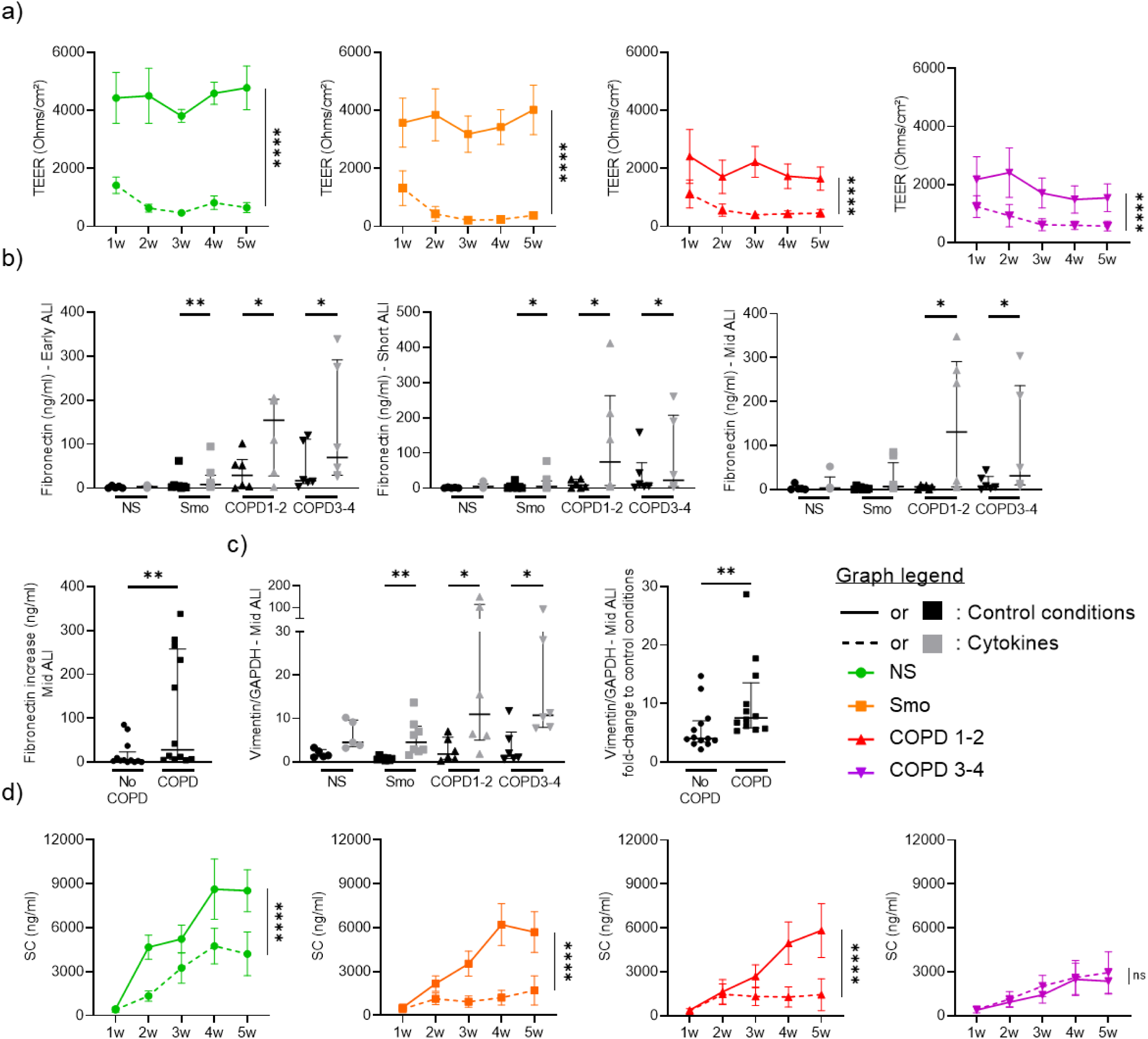
Cytokine activation triggers COPD-like epithelial changes. a) Epithelial inflammation, driven by exogenous TNF-α, IL-1β and IL-6, induces barrier dysfunction, witnessed by a dramatic decrease in TEER in each group (NS, Smo, COPD 1-2, COPD3-4). b-c) Cytokine-induced epithelial inflammation induces EMT in Smo and COPD-derived ALI AE, witnessed by increased fibronectin release in early, short-term and mid-term cultures (b) and vimentin expression in mid-term cultures (c). The absolute increase in fibronectin (b) and vimentin (c) was significantly higher in mid-term COPD AE than in non-COPD AE (pooled Smo and NS, labeled “no COPD”). d) Cytokine-induced epithelial inflammation deteriorates the epithelial polarity, witnessed by the SC apical release. No difference was seen in (very) severe COPD, due to low baseline levels. *, **, **** indicate p-values of less than 0.05, 0.01, and 0.0001. Bars indicate median ± interquartile range (b, c, d) and mean ± SEM (a, e). AE, airway epithelium; ALI, air-liquid interface; COPD, chronic obstructive pulmonary disease; CT, controls; EMT, epithelial-to-mesenchymal transition; GAPDH, glyceraldehyde-3-phosphate dehydrogenase; IL, interleukin; NS, non-smokers; ns, not significant; SC, secretory component; SEM, standard error of the mean; Smo, smokers; TEER, transepithelial electric resistance; TNF-α, tumor necrosis factor α; w, weeks.

In conclusion, exogenous inflammation induces COPD-like changes including barrier dysfunction, EMT and altered polarity in NS- and Smo-AE. In addition, the COPD-derived AE remained prone to develop EMT upon inflammation.

## Discussion

Exploiting unprecedented long-term cultures of ALI-reconstituted human AE from COPD and controls, we demonstrate that the COPD AE retains several abnormalities associated with the disease for prolonged periods of time and reveal key temporal relationships between smoking history and airway cell imprinting as assessed in ALI cultures. We also show that inflammation may recapitulate a COPD-like phenotype and that COPD cells are prone to EMT programming upon inflammatory stimulation.

COPD is a chronic and progressive disease due to repeated injury of the airway epithelial-mesenchymal unit by toxics, leading to AE activation and remodeling. COPD patients who quit smoking may benefit from decreases in mortality, respiratory symptoms and lung function decline (27, 28, 40, 41), and display reduced AE remodeling and goblet cell hyperplasia when smoking cessation exceeds 3.5 years (26). In contrast, no change was observed following smoking cessation regarding axonemal abnormalities in ciliated cells (42), airway mucosal inflammation (30), sputum IL-8/CXCL-8 or neutrophils (29, 43), suggesting permanent alterations in these compartments.

This study explores whether the irreversible nature of the disease is imprinted in the AE in such a way that aberrant features of the native epithelium persist in long-term cultures, independently of signals provided by repeated insults and/or mesenchymal cells and surrounding leukocytes. To study this, we chose to use the ALI model as it was previously shown to recapitulate, at least to some extent, the native COPD phenotype (8, 10, 16, 17, 44), and we prolonged the culture up to 10 weeks, an unusually long and so far unreported duration in this model.

A major abnormality that persists in the COPD AE is the barrier and junctional defect (7), that was initiated in Smo and was associated with decreased protein levels of E-cadherin and occludin (Figures 1&2). This observation corroborates previous studies showing that CS exposure disrupts AJCs (45). We here show that this alteration further persists in long-term cultures for occludin. In addition, changes in TEER and E-cadherin/occludin expression are further aggravated according to the presence and severity of COPD. No difference was observed regarding mRNA abundance for the main AJCs’ components, suggesting post-transcriptional regulation.

In line with the requirement of AJCs’ integrity to ensure baso-apical epithelial polarity (46), we show that AE polarity is persistently impaired in COPD. Aside from abovementioned AJCs disruption, the pIgR/SC system, that allows baso-apical transcytosis of polymeric immunoglobulins (47), was also shown to be defective in COPD (16). Our study demonstrates that this impairment persists over time, with decreased SC apical release and *PIGR* mRNA abundance in COPD3-4 AE (Figure 6). Interestingly, those data are contrasting with previous findings in asthma where pIgR downregulation, although being similarly present *in situ*, does not persist in ALI (48), suggesting distinct mechanisms driving epithelium pathology in asthma and COPD.

Ciliated cell hypoplasia – with decreased *FOXJ1* and *DNAI1* mRNA abundance and reduced bêta-tubulin IV^+^ cells numbers – also persists in long-term COPD-derived cultures, matching *in situ* findings (Figures 3&S3). In addition, goblet cell hyperplasia persists in Smo, as *SPDEF* and *FOXA3* expression, along with MUC5AC^+^ cells numbers, remain higher in active smokers and ex-smokers who quit for less than 4 years, compared with long-quitters (Figure 4). These results support *in situ* data, and corroborate previous findings indicating that the muco-secretory trait relates more directly to smoking rather than to COPD (26).

In contrast, some abnormalities are observed only in short-term cultures. Indeed, we show aberrant EMT features in Smo and COPD AE during the first weeks of culture, with increased fibronectin release and vimentin contents, but these features vanish from mid-term cultures onwards, with complete disappearance occurring earlier in Smo than in COPD (Figure 5). Nevertheless, the COPD AE remains prone to reactivate EMT programming upon inflammatory condition, suggesting the existence of imprinting of the COPD AE by previous (*in vivo*) exposures conditioning its responses to further stimulation (Figure 8).

As observed for EMT, upregulated IL-8/CXCL-8 release by Smo and COPD AE also faded away from mid-term cultures onwards, whereas IL-6 overproduction persisted in long-term ALI-AE from both Smo and COPD patients (Figure 7). *In vitro* exposure to inflammatory cytokines reproduced or aggravated alterations in barrier and polarity features as well as EMT in controls and COPD AE, respectively.

The fundamental mechanisms of these observations question the nature of epithelial memory. Inflammatory memory refers to memories of previous immune events enabling barrier tissues to rapidly recall distinct environmental exposures, which may be stored not only in immune cells but also in epithelial and mesenchymal cells (49, 50). While memory classically refers in the immune system to somatic mutations underlying adaptive antibody responses to recall antigens, “inflammatory memory” is less well defined and may include epigenetic modifications and chromatin changes that may drive persistent alterations in damaged tissues. In the airways, progenitor basal cells are prime candidates to retain epithelial memory, as do epithelium progenitors in the skin towards inflammatory or mechanical stress (50) as well as in the gut towards dietary components (51). Interestingly, WNT signalling, which is upregulated in the COPD AE (17), has been shown to be involved in the latter form of epithelial memory and may regulate stemness and tumorigenicity. In line with the recent hypothesis that basal cells serve as repositories of allergic inflammatory memory in airway epithelial cells (49), one could propose that airway stem cells also store the memory of repeated previous injuries by inhaled toxics such as cigarette smoke.

Our study has several limitations. First, an effect of treatments (e.g., inhaled corticosteroids in severe patients) on the findings cannot be excluded. However, *in vitro* studies on intestinal cell lines and primary HBEC cultured at the ALI showed that dexamethasone induced increased TEER, claudin-2 (52), and E-cadherin (53, 54) expression, while budesonide exposure of ALI-HBEC counter-acted CS-induced barrier dysfunction (55). In addition, dexamethasone and fluticasone propionate improved TGF-β_1_-induced EMT in A549 cells (56), suggesting that corticosteroids may rather improve barrier function and EMT changes. Second, airway progenitor (basal) cells are suspected to display reduced self-renewal (57), although conflicting data exist (58). One cannot exclude that COPD ALI-AE could originate from basal cells with lower replication rate. However, no difference was observed regarding time to reach confluence (both in flasks and in inserts) across study groups. In addition, one could expect a “catch-up effect” in long-term cultures, which was not the case, arguing for a progenitor imprinting in COPD, as already suggested by Ghosh and colleagues (57). Third, the imprinting observed in this model of *ex vivo* reconstituted epithelium that implies proliferation and differentiation of basal cells in prolonged ALI culture, could be confirmed in other models such as 3D airway organoids.

In conclusion, this study demonstrates that the AE from Smo and COPD stores the memory of its native state and previous insults from cigarette smoking. This memory is multidimensional, including alterations in barrier function, epithelial polarity, lineage differentiation, as well as IL-6 release and EMT reprogramming. Further studies are required to explore the mechanisms of this memory, confirm the exact cellular niche retaining it, as well as identify putative targets for future therapies.

## Acknowledgements

The authors would like to thank J. Van Snick for his help with IL-6 ELISA, V. Lacroix, A. Belhaj, Ph. Eucher, and D. Hoton for providing surgical tissue, A. Daumerie for technical advice, M. de Beukelaer and Ch. Fregimilicka for their help with samples processing, and the IREC Pole of Microbiology (UCLouvain, Brussels, Belgium) for sharing their molecular biology facility.

