## Supplementary data for "The memory of airway epithelium damage in smokers and COPD patients"

### Online data supplement

### 1. Supplementary methods

#### Study population and lung tissue samples

Surgical tissue from lobectomies was used from both non-smokers and smokers controls, as well as from mild/moderate COPD patients undergoing lung surgery for a solitary tumor, while lung explants were obtained to analyze lung tissue from patients with (very) severe COPD. Patients with any other lung disease than COPD (e.g., asthma, lung fibrosis) were excluded from the study. All patients received information and signed a written consent to the study protocol, which was approved by the local clinical Ethical committee (reference 2007/19MARS/58 for UCLouvain, S52174 and S55877 for KULeuven).

#### *In vitro* reconstitution of primary human bronchial epithelium on ALI culture

A large piece of lobar or segmentar bronchus (3$^{rd}$ or 4$^{th}$ generation) was selected from lobectomies or explants, located as far as possible from the tumor site (in the case of lobectomies) and submitted to pronase digestion overnight at 4°C, in order to derive primary human broncho-epithelial cells (HBEC) for each sample. HBEC were seeded in 75cm² flasks ($2x10^6$ alive cells/flask), then cultured in retinoic acid-supplemented Bronchial Epithelial Cell Growth Basal Medium (BEBM, Lonza, Verviers, Belgium) until confluence, which was reached at day 7 (+/- 2 days [SD]), with no difference across the groups. Confluent cells were exclusively (> 99,9%) $p63^+$ basal cells. Cells were then detached (passage 1), assessed for viability and seeded at a density of 80,000 alive cells/well on 24-well polyester filter-type inserts (0.4-µm pore size; Corning, Corning, NY) coated with 0.2 mg/ml collagen IV (Sigma-Aldrich, Saint-Louis, MO) until a confluent monolayer was obtained. Of note, mean cell viability before seeding in inserts was 92,4% (+/- 3,4% [SD]), and mean time to confluence in submerged conditions was 6 days (+/- 0.8 day [SD]), with no difference between the study groups. The culture was then carried out in ALI.

Twenty-six samples were carried out for 10 weeks and assessed for several epithelial properties. Twenty-five samples were carried out for 5 weeks and used exclusively for the 'inflammatory stimulation' experiment (see details below). Once in ALI, HBEC were cultured in BEBM:Dulbecco's Modified Eagles Medium (DMEM) (1:1) medium supplemented with penicillin (100U/ml), streptomycin (100µg/ml) (Lonza, Verviers, Belgium), bovine serum albumin (BSA) (1.5 µg/ml), retinoic acid (30ng/ml) (Sigma, Saint-Louis, MO), and BEGM SingleQuotsTM Supplements and Growth Factors (Lonza), including bovine pituitary extract (52µg/ml), insulin (5µg/ml), hydrocortisone (0.5g/ml), transferrin (10µg/ml), epinephrine (0.5µg/ml), epidermal growth factor (0.5ng/ml) and triiodothyronine (3.25ng/ml). No culture infection occurred among the 51 samples.

Every week during ALI culture, basolateral media were collected, and the apical pole of HBEC was washed with 300µl sterile phosphate-buffered saline (PBS) before centrifugation for 5 minutes at 10,000g. Transwell inserts were fixed by direct immersion in 4% buffered formaldehyde, before incubation in PBS (pH 7.4) and embedding in paraffin blocks. ALI-HBEC were also processed for mRNA abundance or Western blot analyses (see below).

The 25 samples that underwent 5 weeks ALI culture were exposed (48h) or not (control condition) to a pro-inflammatory cytokine cocktail including IL-1β, IL-6, TNF-α, each at 5 ng/ml (Miltenyi Biotec, Germany) in the basolateral compartment following a preliminary titration experiment (10, 5, and 2,5 ng/ml) where no cytotoxicity (release of lactate dehydrogenase < 5%) was shown at 5 ng/ml (data not shown).

**Reverse transcriptase quantitative polymerase chain reaction (RT-qPCR)**

Total RNA was extracted from reconstituted ALI cultured epithelia using TRIzol reagent (Thermo Fisher Scientific). 500ng of RNA was reverse-transcribed with RevertAid H minus Reverse transcriptase kit with 0.3 µg of random hexamer, 20U of RNase inhibitor and 1mM of each dNTP (Thermo Fisher Scientific, Waltham, MA) following the manufacturer's protocol in a thermocycler (Applied Biosystems, Foster City, CA). The expression levels were quantified by real-time quantitative PCR with the CFX96 PCR (Bio-Rad, Hercules, CA). The reaction mix contained 2.5 µl of complementary desoxyribonucleic acid diluted 10-fold, 200nM of each primer (primers properties are detailed in Table S2) and 2x iTaq UniverSybr Green® Supermix (Bio-Rad) in a final volume of 20 µl. The cycling conditions were 95°C for 3 minutes followed by 40 cycles of 95°C for 5 seconds and 60°C for 30 seconds. To control the specificity of the amplification products, a melting curve analysis was performed. The copy number was calculated from the standard curve. Data analysis was performed using Bio-Rad CFX software (Bio-Rad). Expression levels of target genes were normalized to the geometric mean of the values of 3 housekeeping genes (RPL27, RPS13, RPS18).

**Western blot assays**

Cells were lyzed with 150µl of Laemmli's sample buffer containing 0.7M 2-mercaptoethanol (Sigma-Aldrich) and lysates were stored at -20°C. After thawing, samples were heated at 100°C for 5 minutes, loaded in a SDS-PAGE gel before migration at 100V for 15 minutes and then at 180V for 50 minutes. Cell proteins were transferred onto a nitrocellulose membrane (Thermo Fisher Scientific) at 0.3A for 2 hours 10 minutes at RT. The membranes were blocked with 5% w/v BSA (Sigma-Aldrich) in Tris-buffered saline with 0.1% Tween 20 (Sigma-Aldrich) for 1 hour at RT, then washed and incubated overnight at 4°C with a primary antibody according to the target protein (see Table S3 listing used primary and secondary antibodies). Membranes were then incubated for 1 hour at room temperature (RT) with horseradish peroxidase (HRP)-conjugated secondary anti-rabbit (Cell Signalling, Danvers, MA) or anti-mouse (Sigma) IgG.

**Sandwich enzyme-linked immunosorbent assay (ELISA) for SC, IL-8/CXCL-8 and IL-6**

Basolateral IL-8/CXCL8 and IL-6 release were assessed by sandwich ELISA, following manufacturer's instructions (of IL-6). Briefly, 96-well plates were coated overnight, at 4°C, with anti-IL-8/CXCL8, -IL-6, and -SC antibodies diluted in bicarbonate buffer (pH 9,6). Then, after blocking with 1% w/v BSA in phosphate-buffered saline for 90 min at 37°C, HBEC apical washes (for SC) or basolateral supernatants (for IL-8/CXCL-8 and IL-6) were incubated for 60 min at 37°C, along with standard samples. Detection was performed with a first incubation with the corresponding biotinylated antibody (anti-fibronectin, -SC, -IL-8/CXCL-8 or -IL-6), followed by a second incubation with HRP-linked anti-mouse IgG, for 1 h each. Revelation was performed with 3,3',5,5'-tetramethylbenzidine (TMB, Fisher) and stopped with $H_2SO_4$ 1.8 M.

**Direct ELISA for fibronectin**

HBEC basolateral washes and fibronectin standard were coated in plates with bicarbonate buffer overnight (pH 9,6), at 4°C. After blocking with 1% w/v BSA in phosphate-buffered saline for 90 min at 37°C, detection was performed with a first incubation with mouse anti-fibronectin, followed by a second incubation with HRP-linked anti-mouse IgG, for 1 h each. Revelation was performed with TMB and stopped with $H_2SO_4$ 1.8 M.

**Immunofluorescence staining using tyramide signal amplification**

Five micron-sections of bronchial tissue or reconstituted ALI epithelium, fixed in 4% formaldehyde and paraffin-embedded, were deparaffinised in toluene and rehydrated through a graded series from ethanol to water. Antigen retrieval was performed in citrate buffer (pH 6.0 containing 0.1% of triton) using a pressure cooker at 15 PSI for 5 minutes. Sections were blocked for non-specific antigen binding by incubation in Bloxall (Vector Laboratories Inc.) for 15 min and then in 0.3% hydrogen peroxide with 5% goat serum (Bio-Rad) for 30 min. Staining first included a 30 min protein blocking with 5% goat serum, then the primary antibodies diluted in 5% normal goat serum solution were applied, then the appropriate SuperBoost™ goat anti-rabbit or anti-mouse, poly-HRP-conjugated secondary antibody (Thermo Fisher Scientific) was applied for 40 min. HRP-conjugated polymer mediated the focal covalent binding of a fluorophore using tyramide signal amplification. In surgical samples, this cycle was repeated twice to allow multiplex staining of pIgR, bêta-tubulin IV, and MUC5AC. Table S4 recapitulates the antibodies and fluorophores that were used. Finally, sections were counterstained with Hoechst (Thermo Fisher Scientific) diluted at 10µg/ml in TBS-BSA 5% and mounted with Dako fluorescence mounting medium (Dako, Carpinteria, CA). For negative controls, we used rabbit or mouse isotype controls at the same concentration as the corresponding primary antibodies (diluted in 5% normal goat serum).

### Immunofluorescence staining quantification

For ALI-reconstitued AE, bêta-tubulin IV$^+$ and MUC5AC$^+$ cells were manually counted at different time-points (1 week, early; 2 weeks, short-term; 4 weeks, mid-term; and 9 weeks, long-term). For each sample, 5 fields (20x magnification) were analysed, and the arithmetic mean of the 5 fields was calculated.

For *in situ* bronchial sections, QuPath 0.4.2 analysis tool (1) was used. First, epithelial layers were manually delineated on each slide. Staining thresholds for MUC5AC and bêta-tubulin IV staining were defined prior to software analysis. Finally, the positive surface was calculated within the total area delineated and expressed as a percentage.

### Statistical analysis

Prior to analysis, all data were assessed for normality using Shapiro-Wilk and Kolmogorov-Smirnov tests. Parametric tests were used only when all groups were considered normal with the two tests. Data were expressed as means and standard deviation for data reaching normality, while non-normally distributed data were expressed as medians and interquartile ranges. Data were analysed with JMP® Pro, Version 14 (SAS Institute Inc., Cary, NC, USA) and GraphPad Prism version 8.0.2 for Windows (GraphPad Software, La Jolla, CA, USA). p-values $< 0.05$ were considered statistically significant.

For timepoint analyses, Brown-Forsythe and Welch ANOVA tests, followed by Holm-Sidak's multiple comparisons tests (each experimental group versus non-smoker controls) were used for normally distributed data (Figures 1a, 2b, 3a&c, 4b&c, 6a,c&e, 7a) while Kruskal Wallis tests followed by Dunn's multiple comparisons tests (each experimental group versus non-smoker controls) were used for non-normally used data (Figures 2a, 3b&e, 4a, 5a&c, 6d, 7b).

For two-groups comparisons, unpaired t-tests were used for normally distributed data (Figure 4c [right graph]), while Mann-Whitney U tests were used for non-normally distributed data (Figure 4d). For paired data (Figure 8), paired Wilcoxon signed rank tests were used (Figures 8b and 8c)

All correlations were assessed by using linear regression and Cohen's kappa coefficient calculation.

For longitudinal analyses (Figures 1b, 3a-b [middle graphs], 4a [right graph], 5c [right graph], 6a&c [right graphs], 7a-b[right graphs] and 8a&d), linear mixed models were built using JMP® Pro, Version 14. The models aimed at examining the effects of time (weeks of culture) and phenotype of the sample on continuous variables, and were designed to take into account repeated measures (i.e., measurements taken at different timepoints on the same samples), and specified a nesting structure where each sample was nested within a study group (i.e. NS, Smo, COPD1-2, COPD3-4). A random effect for each nested sample "sample[phenotype]" was also integrated before the model was run. Post-hoc analysis with a correction of least significant difference (LSD) for multiple comparisons was performed between the main effects.

### 2. Supplementary tables

| Total n=26 | Non-smoker controls (n=5) | Smoker controls (n=8) | COPD1-2 (n=6) | COPD3-4 (n=6) | |
|---|---|---|---|---|---|
| **N (Male/Female)** | 5 (2/3) | 8 (5/3) | 6 (2/4) | 6 (3/3) | ns |
| **Age** | 69.4 ± 14.1 | 62.4 ± 7.1 | 62.0 ± 4.6 | 61.7 ± 2.1 | ns |
| **Smoking history (never/former/current n)** | 5/0/0 | 0/5/3 | 0/3/3 | 0/6/0 | p<0.01 |
| **Pack-years** | NA | 37.0 ± 27.1 | 42.7 ± 24.6 | 53.3 ± 30.3 | ns |
| **If applicable, duration since smoking cessation (months)** | NA | 275.4 ± 149.7 | 85.6 ± 56.9 | 108.1 ± 116.5 | ns |
| **FEV1 (% of PV)** | 110.8 ± 18.5 | 99.1 ± 9.6 | $70.3 \pm 8.0^{*\#}$ | $26.7 \pm 7.8^{*\#¶}$ | p<0.0001 |
| **FEV1/VC ratio (%P)** | 79.7 ± 8.7 | 74.3 ± 3.3 | $66.0 \pm 7.9^{*\#}$ | $35.2 \pm 7.5^{*\#¶}$ | p<0.0001 |
| **DLCO (% of PV)** | 92.8 ± 9.9 | 88.5 ± 10.7 | 64.2 ± 15.3 | $36.5 \pm 8.3^{*\#}$ | p<0.01 |
| **BMI (kg.m$^{-2}$)** | 26.7 ± 7.6 | 26.7 ± 5.3 | 27.5 ± 6.2 | 25.4 ± 4.2 | ns |
| **Inhaled corticosteroids (n/total N)** | 0/5 | 0/8 | 1/6 | 5/6 | p<0.001 |
| **Surgical indication** | | | | | |
| - **Neoplasia** | 4/5 | 8/8 | 5/6 | 0/6 | |
| ○ **SCC** | 0 | 2 | 1 | 0 | |
| ○ **AC** | 2 | 4 | 3 | 0 | |
| ○ **Carcinoid tumour** | 2 | 1 | 0 | 0 | |
| ○ **Pulmonary metastasis of other cancers** | 0 | 1 | 1 | 0 | |
| - **Lung transplant** | 0 | 0 | 0 | 6 | |
| - **Other** | 0 | 0 | 1 | 0 | |
| - **Declined lung donor** | 1 | 0 | 0 | 0 | |

**Table S1 | Patient series for short-term (5 weeks) ALI culture with/without inflammatory condition.** Data are presented as mean ± SD, unless otherwise stated. Demographic data, lung function tests, smoking history and inhaled corticotherapy are stated for the patient groups, classified according to smoking history and the presence and severity of airflow limitation. AC, adenocarcinoma; ALI, air/liquid interface; BMI, body mass index; COPD, chronic obstructive pulmonary disease; DLCO, diffusing capacity of the lung for CO; FEV1, forced expiratory volume in 1 s; FVC, forced vital capacity; NA, not applicable; PV, predicted values; SCC, squamous cell cancer.
* = p<0.05 compared to non-smoker controls.
$^{\#}$ = p<0.05 compared to smoker controls
$^{¶}$ = p<0.05 compared to COPD1-2 patients
ns, not significant

| Gene | FORWARD primer (5'-3') | REVERSE primer (3'-5') | Amplicon size (bp) | Melting t° (°C) |
|---|---|---|---|---|
| **Housekeeping genes** | | | | |
| *RPL27* | TGG TAG GGC CGG GTG GTT GC | ACT TTG CGG GGG TAG CGG TC | 185 | 60 |
| *RPS13* | TCG GCT TTA CCC TAT CGA CGC AG | ACG TAC TTG TGC AAC ACC ATG TGA | 153 | 60 |
| *RPS18* | TGT GGG CCG AAG ATA TGC T | TGA TCA CAC GTT CCA CCT CAT | 101 | 60 |
| **Target genes** | | | | |
| *CDH1* | GCT GGA CCG AGA GAG TTT CC | CGA CGT TAG CCT CGT TCT CA | 179 | 60 |
| *CXCL8* | CTC TGT GTG AAG GTG CAG TTT TG | AAC TTC TCC ACA ACC CTC TGC | 223 | 60 |
| *DNAI1* | GAG TTG ACC GAT GCG GAG TT | GGT TCC CAA CCT GGG TGT AG | 163 | 60 |
| *DNAI2* | GCG ATT CAT ACA TCT GGG AC | CAG CAG GCT ATC TGT CCA T | 145 | 60 |
| *FOXA3* | TGC TGG GCT CAG TGA AGA TG | GTC ATG TAG GAG TTG AGG GGG | 124 | 60 |
| *FOXJ1* | CCT GGC AGA ATT CAA TCC G | GCG TAC TGG GGG TCA AT | 115 | 60 |
| *IL6* | AGT TCC TGC AGA AAA AGG CAA AG | TGA GGT GCC CAT GCT ACA TTT | 198 | 60 |
| *MCIDAS* | GAC GCG CTT GTT GAG AAT AA | CAC GTT CCG CTC CTT GAG | 84 | 60 |
| *MYB* | TAC TGC CTG GAC GAA CTG ATA A | CTG GCT GGC TGG CTT TTG AA | 111 | 60 |
| *PIGR* | CTC TCT GGA GGA CCA CCG T | CAG CCG TGA CAT TCC CTG | 78 | 60 |
| *SPDEF* | TGA CCT TGG AGG AGC ACT CG | CAT GGG ATC TGC GGT GAT GTT | 110 | 60 |
| *TJP1* | GTG GTT CTT CGA GAA GCT GGA | TGC AGG CGA ATA ATG CCA GA | 170 | 60 |
| *VIM* | CGG GAG AAA TTG CAG GAG GA | AAG GTC AAG ACG TGC CAG AG | 105 | 60 |

**Table S2 |** List of the primers used for RT-qPCR.

| Target | Species | Brand and reference |
|---|---|---|
| **Primary antibodies** | | |
| *E-cadherin* | Mouse monoclonal antibody | Dako M3612 |
| *GAPDH* | Rabbit polyclonal antibody | Sigma G9545 |
| *pIgR* | Rabbit polyclonal antibody | Home made |
| *Occludin* | Rabbit polyclonal antibody | Merck-Millipore ABT146 |
| *Vimentin* | Mouse monoclonal antibody | Dako M0725 |
| **Secondary antibodies** | | |
| *Anti-Rabbit HRP-conjugated* | Goat polyclonal antibody | Cell signalling 7074S |
| *Anti-mouse HRP-conjugated* | Sheep polyclonal antibody | Sigma A6782 |

**Table S3 |** List of the primary and secondary antibodies used for western blot.

| Target | Species | Brand and reference |
|---|---|---|
| **Primary antibodies** | | |
| *bêta-tubulin IV* | Mouse monoclonal antibody | Sigma T7941 |
| *MUC5AC* | Mouse monoclonal antibody | Acris AM50143PU-N |
| *pIgR* | Rabbit polyclonal antibody | Home made |
| *Mouse* $IgG_1$ *Isotype* | Mouse monoclonal antibody | eBiosciences 14-4714-82 |
| *Rabbit IgG Isotype* | Rabbit polyclonal antibody | Homemade antibody |
| **Secondary antibodies** | | |
| *Anti-Rabbit HRP-conjugated* | Goat polyclonal antibody | Cell signalling 7074S |
| *Anti-mouse HRP-conjugated* | Sheep polyclonal antibody | Sigma A6782 |

**Table S4 |** List of the primary and secondary antibodies used for tyramide signal amplification-enhanced immunofluorescence staining.

### 3. Supplementary Figures

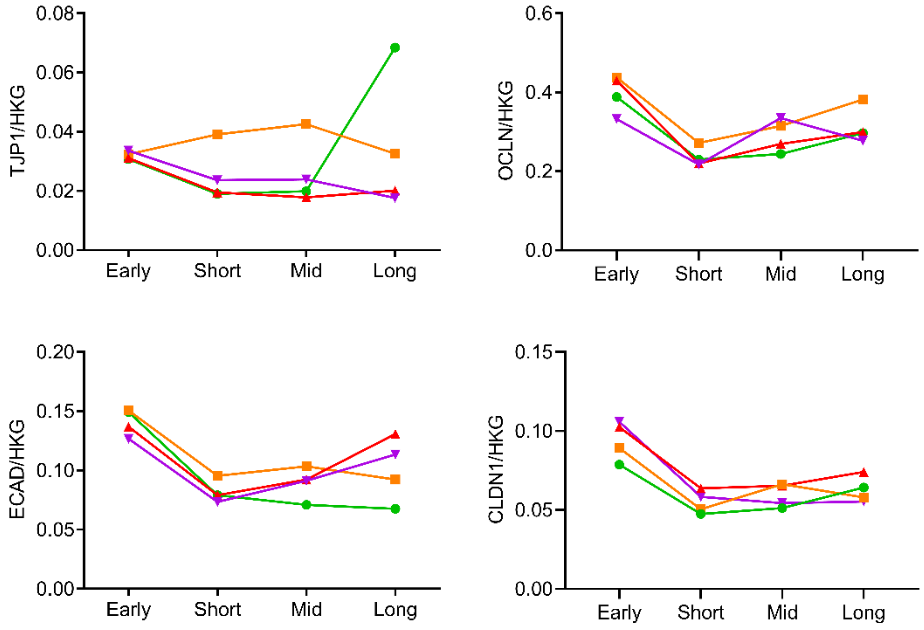


**Figure S1 | Evolution over time of the gene expression of key components of tight and adherens junctions of the AE**

No difference was seen over time regarding the mRNA abundance of *TJP1*, *OCLN*, *ECAD* and *CLDN1* in the AE of non-smokers, smoker controls, mild-to-moderate COPD and (very) severe COPD patients.

Graphs plot means. No error bar are provided to keep the graphs readable. AE, airway epithelium; COPD, chronic obstructive pulmonary disease; NS, non-smokers; Smo, smokers; PV, predicted values.

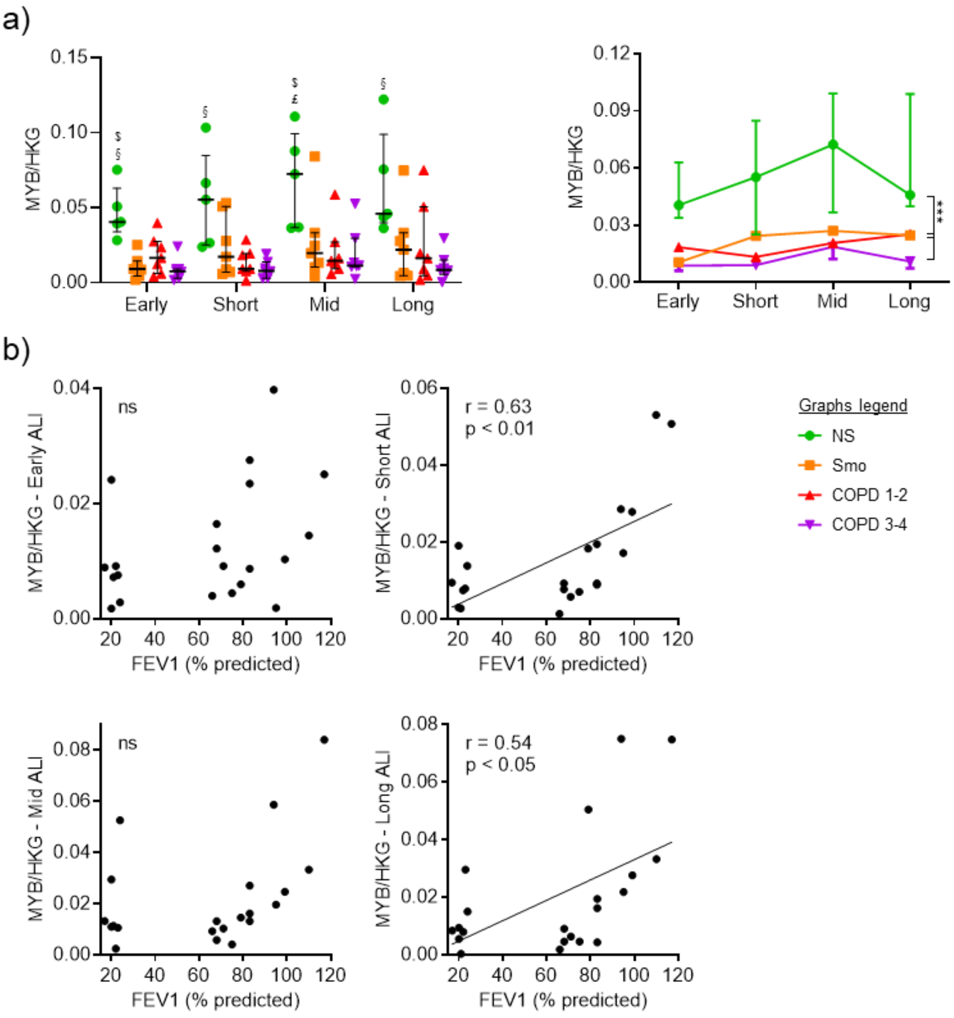


**Figure S2 | Altered differentiation towards pre-differentiated cells in the COPD AE.**
a) Decreased *MYB* expression in early, short-term, mid-term and long-term ALI culture in smokers and COPD AE, as compared with non-smokers.
b) In short- and long-term cultures, *MYB* mRNA levels downregulation correlated moderately but significantly with the COPD severity, witnessed by the FEV1. Correlation graph includes only Smo and COPD samples to specifically assess the correlation with the disease severity.
$, £ and § highlight significant decreases in Smo, COPD1-2 and COPD3-4 patients respectively, as compared to NS. ** indicates p-values of less than 0.01. Bars indicate median ± interquartile range
AE, airway epithelium; ALI, air-liquid interface; COPD, chronic obstructive pulmonary disease; CT, control; FEV1, forced expired volume in 1 second; HKG, housekeeping genes; NS, non-smokers; Smo, smokers; y, years.

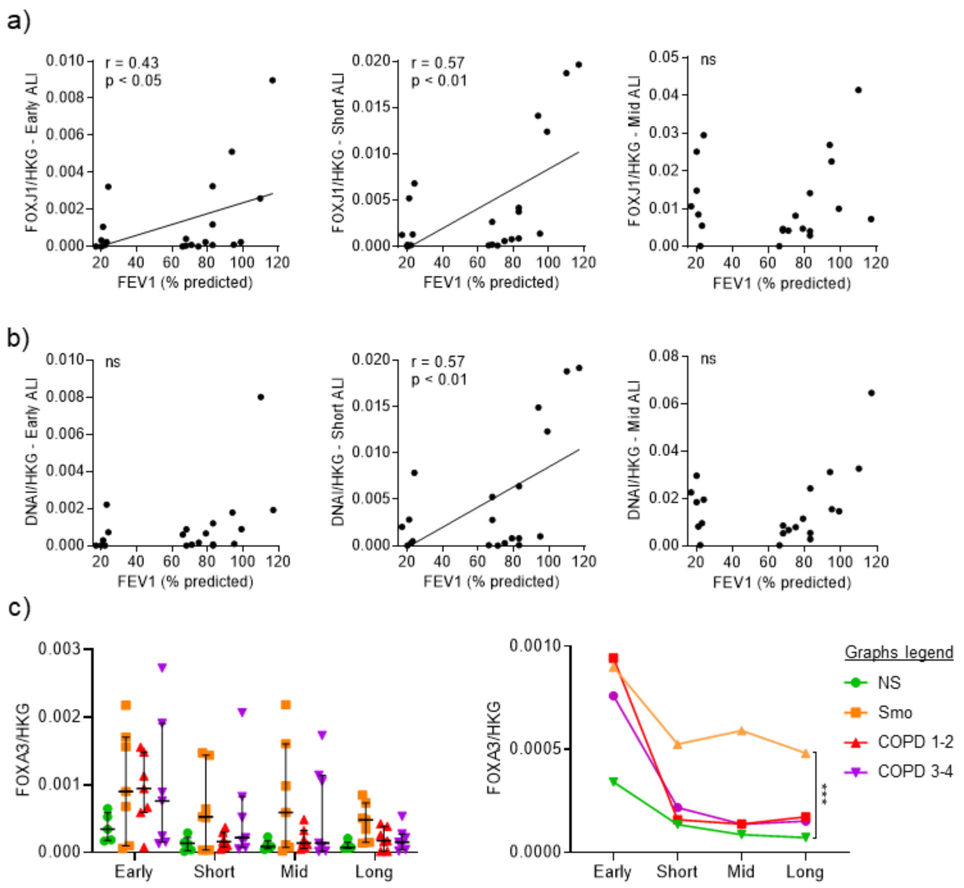


**Figure S3 | Altered differentiation towards ciliated and goblet cells, in COPD and smokers airway epithelium, respectively**

a) In early and short-term cultures at the ALI, COPD-related *FOXJ1* downregulation correlates with the disease severity, witnessed by the FEV1. Correlation graphs include only Smo and COPD samples to specifically assess the correlation with the disease severity.

b) In short-term cultures at the ALI, COPD-related *DNAI1* downregulation correlates with the disease severity, witnessed by the FEV1. Correlation graphs include only Smo and COPD samples to specifically assess the correlation with the disease severity.

c) *Left panel*: *FOXA3* mRNA level in Smo ALI-AE is non-significantly increased at all time periods, as compared with non-smokers. *Right panel*: Longitudinal analysis of *FOXA3* mRNA levels, confirming the upward trend and depicting significantly increased *FOXA3* expression in Smo AE as compared with NS.

*** indicates p-values of less than 0.001. Bars indicate median ± interquartile range, except for g, mean ± SEM. AE, airway epithelium; ALI, air-liquid interface; COPD, chronic obstructive pulmonary disease; FEV1, forced expired volume in 1 second; NS, non-smokers; ns, not significant; SEM, standard error of the mean; Smo, smokers; PV, predicted values.

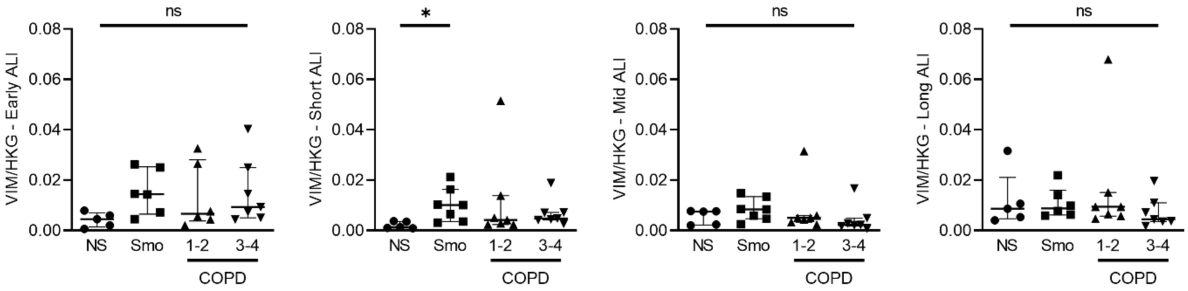


**Figure S4 | Vimentin mRNA levels do not vary in Smo and COPD AE as compared with NS AE.** No difference was observed regarding *VIM* expression in smokers and COPD-derived AE, as compared with that of NS, at all time periods, except for an increased expression in Smo in short-term ALI cultures. * indicates a p-value of less than 0.05 (analyzed using the Kruskal-Wallis test followed by Dunn's post-hoc test). Bars indicate median ± interquartile range. AE, airway epithelium; ALI, air-liquid interface; COPD, chronic obstructive pulmonary disease; NS, non-smokers; ns, not significant; Smo, smokers.

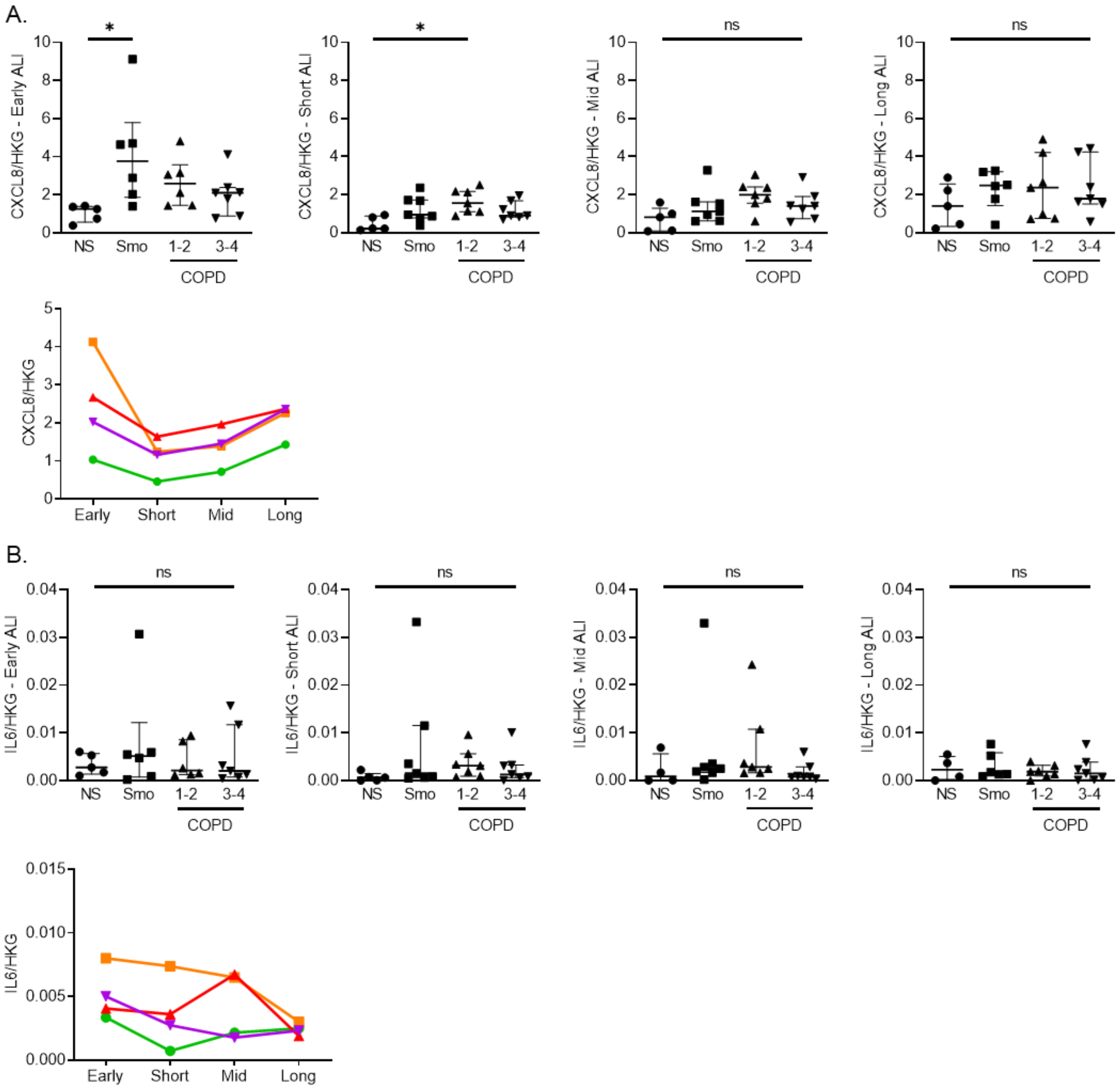


**Figure S5 | *IL8/CXCL8* and *IL6* mRNA abundance is not significantly modified in smokers and COPD AE.**
**A.** *IL8/CXCL8* gene expression was enhanced in smokers and mild-to-moderate COPD in early and short-term ALI cultures, respectively, but no striking difference was observed in later time periods, nor in (very) severe COPD.
**B.** No difference was observed in *IL6* gene expression in smokers and COPD-derived ALI-AE as compared with non-smokers.
* indicates a p-value of less than 0.05 (analyzed using the Kruskal-Wallis test followed by Dunn's post-hoc test). Bars indicate median ± interquartile range, except for longitudinal analysis, mean. AE, airway epithelium; ALI, air-liquid interface; COPD, chronic obstructive pulmonary disease; NS, non-smokers; ns, not significant; SEM, standard error of the mean; Smo, smokers; transepithelial electric resistance.

**Supplementary data references**

1. Bankhead P, Loughrey MB, Fernandez JA, Dombrowski Y, McArt DG, Dunne PD, McQuaid S, Gray RT, Murray LJ, Coleman HG, James JA, Salto-Tellez M, Hamilton PW. QuPath: Open source software for digital pathology image analysis. *Sci Rep* 2017; 7: 16878.
